# Comparison and Evaluation of Neutralization of Clinically Frequently Used Antimicrobial Agents Using Three Different Culture Media in Simulated Blood Cultures

**DOI:** 10.1101/2024.01.15.575683

**Authors:** Yaping Hang, Jianqiu Xiong, Longhua Hu, Yanhui Chen, Shan Zou, Xueyao Fang, Yanping Xiao, Xingwei Cao, Hong Lou, Xiuzhen Li, Yanhua Liu, Qiaoshi Zhong

**Affiliations:** Department of clinical laboratory, the 2^nd^ affiliated Hospital, Jiangxi Medical College, Nanchang University.Nanchang; 330006 P.R China; Intravenous medication dispensing center, the 2^nd^ affiliated Hospital, Jiangxi Medical College, Nanchang University.Nanchang; 330006 P.R China

**Keywords:** BACT/ALERT plus media, DL media, Thermo media, blood culture, antimicrobial neutralization, recovery rate, aerobic, anaerobic

## Abstract

The performance of BACT/ALERT FA/FN Plu (France) blood culture containing a novel resin, DL (China) blood culture containing common resin, and adsorbent-free Thermo (USA) blood culture relying on dilution for antimicrobial neutralization at %peak serum concentration was evaluated by measuring the recovery of organisms and time to detection (TTD) in nine simulated microorganism-antimicrobial combination blood cultures.There was a significant difference in recovery rates between the aerobic media: 35/40 (87.5%) for BACT/ALERT media, 15/35 (42.9%) for DL media, and 5/40 (12.5%) for Thermo media; whereas, there was no statistical difference in TTD between FA Plus media and DL aerobic media. The recovery rates of the anaerobic media were 32/35 (91.4%) for BACT/ALERT media, 1/35 (2.9%) for DL media, and 5/35 (14.3%) for Thermo media, with significant differences only between BACT/ALERT FN Plus media and the others. Among the seven main antimicrobial categories, only BACT/ALERT FA/FN Plus culture media showed high recovery of microorganisms, except for carbapenems. The DL culture media exhibited a relatively high recovery rate of piperacillin/tazobactam, levofloxacin, and gentamicin only in aerobic media, and Thermo media only displayed the recovery of gentamicin.BACT/ALERT FA/FN Plus culture media with novel resin showed absolute advantages over DL and Thermo culture media and, thus, can be selectively applied clinically with prior use of antimicrobials before pathogen detection. DL culture media containing common resin outperformed adsorbent-free dilution-based Thermo culture media, which can be a backup option. We should focus on improving the neutralization of carbapenems with low efficiency in all three media.

## Introduction

Accurate and timely diagnosis of bloodstream infections (BSIs) is crucial for the proper management of infected patients, particularly those with infection-complicated sepsis and septic shock, and for reducing the associated high morbidity and mortality (1). Blood cultures remain the gold standard for BSI detection and should be collected when sepsis is suspected in patients before antimicrobial drug administration. However, in an urgent sepsis condition, the supervising physician empirically provides broad-spectrum antimicrobial therapy before obtaining a microbiological diagnosis of the infection (2). To minimize the inhibitory effect of antimicrobials on microbial growth in blood culture, certain commercial blood culture media contain substances, including resin or charcoal, that are intended to adsorb antimicrobials or other substances and have been developed to improve the detection of microorganisms from sepsis patient samples (3–4). Most studies have primarily focused on the capacity of bacterial pathogenic detection in blood cultures. Some recent blood culture simulation studies (involving blood culture media injected with whole blood, antimicrobials, and microorganisms) have assessed the antimicrobial-neutralizing capability of various blood culture media from different manufacturers with different mechanisms (5–6).

This study aimed to compare the neutralization effects of three automated blood culture systems that utilize three manufacturer blood culture media with different antibacterial mechanisms, including recently approved China mainland BACT/ALERT (FA Plus and FN Plus) media (bioMérieux, France), widely used in China mainland DL (ZhuhaiDL, China), and Thermo blood culture media (Thermo Scientific, USA).

## Materials and methods

### Ethics Clarification

No ethical review process was required for this simulated blood culture study because it did not involve any animal or human experiments. Moreover, as a simulated study, no specimens, microbial isolates, or any other material from patients or healthy human bodies were involved. All materials used in this study, including reference ATCC microorganism strains, antimicrobial agents, and horse blood, were commercially available and purchased.

### Blood culture media and instruments

In this study, 40 mL of ZLAPB resin-containing BACT/ALERT FA Plus aerobic, BACT/ALERT FN Plus anaerobic blood culture media, and 40 mL of adsorbent-free BacT/ALERT SA aerobic (SA) and SN anaerobic (SN) media were applied to the automated BACT/ALERT 3D blood culture system. Moreover, 30 mL of resin-containing DL aerobic and DL anaerobic (commonly used in China) blood culture media was combined with an automated DL-Bt blood culture system. Furthermore, 80 mL of adsorbent-free Thermo VersaTREK REDOX 1 (Thermo Aerobic) and Thermo VersaTREK REDOX 2 (Thermo Anaerobic) blood culture media was tested using an automated Thermo Scientific™ VersaTREK™ blood culture system.

### Microorganisms and antimicrobial substances

Microbial species frequently isolated in clinical microbiology laboratories were selected. The reference strains used included methicillin-susceptible *Staphylococcus aureus* ATCC 29213, *Streptococcus pneumoniae* ATCC 49619, *Escherichia coli* ATCC 25922, *Pseudomonas aeruginosa* ATCC 27853, *Candida albicans* ATCC 90028, and *Bacteroides fragilis* ATCC 25285. The corresponding clinical strains sensitive to the antimicrobial agents used in this study were also tested. Colonies from Columbia blood agar plates were suspended in physiological saline and serially diluted to a 10^2^ CFU/mL target suspension. Concurrently, 0.3 mL of the final dilutions were plated on Columbia blood agar plates and incubated at 37 °C overnight to validate the CFUs.

The nine most commonly used antimicrobials for treating specific BSIs were selected (Table 1). Commercially available standards of penicillin G, piperacillin-tazobactam, meropenem, imipenem, gentamicin, cefoxitin, levofloxacin, vancomycin, and fluconazole (Meilunbio, China) were dissolved in a suitable solvent according to the CLSI guidelines and the manufacturer’s instructions. Each antimicrobial agent was diluted in sterile water or PBS from stock solutions to ensure that 0.5 mL of the dilution contained the desired final drug concentration for each culture media. Antimicrobial solutions at peak serum concentrations (%PSL) achieved after standard adult dosing were used to simulate patient blood levels according to CLSI M100, M60, CNAS-GL028, and the Sanford Guide to Antimicrobial Therapy (50^th^ Edition; Table 1).

**Table 1.** Microorganism and Antimicrobial Combination Tested by the Simulated Adult Blood Culture Model.

**Note:**<sup>a</sup>MICs of the antimicrobial drugs for each strain tested were determined by CLSI broth microdilution and susceptibility categorization was assigned based on published CLSI breakpoints, S: susceptible.
| Species(Drug) | Antibiotic | Species (Strain) | Atmosphere |  | MIC<sup>a</sup><br>(µg/mL) | PSL<sup>b</sup><br>(µg/mL) | %PSL<sup>c</sup><br>(µg/mL) |
| --- | --- | --- | --- | --- | --- | --- | --- |
|  |  |  | Aerobic | Anaerobic |  |  |  |
| Penicillins | Penicillin G | *Streptococcus pneumoniae*(ATCC49619) | √ | √ | 0.25-1(S) | 20 | 100 |
|  | Piperacillin/<br>tazobactam | *Pseudomonas aeruginosa*(ATCC27853) | √ |  | 1/4-8/4(S) | 242/24 | 100 |
| Carbapenems | Meropenem | *Bacteroides fragilis*(ATCC25285) |  | √ | 0.03-0.25(S) | 49 | 100 |
|  | Imipenem | *Escherichia coli* (ATCC25922) | √ | √ | 0.06-0.25(S) | 40 | 100 |
| Aminoglycosides | Gentamicin | *Escherichia coli* (ATCC25922) | √ | √ | 0.25-1(S) | 10 | 100 |
| Cephameycins | Cefoxitin | *Staphylococcus aureus*(ATCC29213) | √ | √ | 1-4(S) | 110 | 100 |
| Quinolones | Levofloxacin | *Staphylococcus aureus*(ATCC29213) | √ | √ | 0.06-0.5(S) | 8.6 | 100 |
| Glycopeptides | Vancomycin | *Staphylococcus aureus*(ATCC29213) | √ | √ | 0.5-2(S) | 50 | 100 |
| Triazoles | Fluconazole | *Candida albicans*(ATCC90028) | √ |  | 0.25-1(S) | 14 | 100 |

### Blood culture

Blood culture media were inoculated with 9 mL of phosphate-buffered saline (PBS) or sterile horse blood (Oxoid), 0.3 mL of microorganism suspension (30-10^2^ CFU/bottle), and 0.5 mL of antimicrobial solution at %PSL. Microorganisms were simultaneously spiked into aerobic and anaerobic media as pairs, except for *P. aeruginosa* and *C. albicans*, which were only evaluated under aerobic conditions, and *B. fragilis*, tested under anaerobic conditions. Considering the lower neutralization rates with carbapenems (5), incubations were performed under anaerobic conditions with carbapenems.

For each microorganism-antimicrobial combination, incubations were performed five times for antimicrobial neutralization tests of three different blood culture media and in triplicate for contrast tests. To ensure the efficacy and susceptibility of the tested antimicrobial, positive controls with 9 mL PBS or horse blood, 0.3 mL bacterial suspension, and 0.5 mL antimicrobial solution at % PSL were incubated in adsorbent-free SA/SN standard media. Simultaneously, to ensure the activity of the ATCC strains, positive controls with 9 mL PBS or horse blood and 0.3 mL bacterial suspension were incubated in every blood culture medium. Additionally, negative antimicrobial-free controls (containing only PBS or horse blood in SA/SN bottles) were used to confirm the sterility of PBS or horse blood. After five days, all culture media were incubated until flagged as positive or negative.

### Data analysis and statistics

Statistical analyses were conducted using GraphPad Prism 7. The detection rates in aerobic and anaerobic media between the three culture systems were compared using Fisher’s exact test. Mann-Whitney tests were applied to analyze the variations in time to detection (TTD), and the median of TTD was compared because the data were not normally distributed. While calculating the median TTD, the results from microorganisms that remained undetected within five days were excluded. A *p-*value of < 0.05 was considered statistically significant.

## Results

In the BACT/ALERT 3D blood culture system, 40 FA Plus media and 35 FN Plus media were tested. In the DL-Bt blood culture system, 35 DL aerobic and 35 DL Anaerobic media were examined, and in the Thermo Scientific™ VersaTREK™ blood culture system, 40 Thermo aerobic and 35 Thermo anaerobic media were evaluated. All microorganisms tested in the antimicrobial-free control media were recovered, except for *C. albicans* in the DL aerobic media (97.8%; 132/135). Moreover, *Candida. spp*. exhibited poor growth in anaerobic media, and positive controls of recovering *C. albicans* in DL aerobic media were negative. Therefore, the *C. albicans*-fluconazole combination in DL media was excluded. When antibiotics were present, eight of the nine antimicrobial agents exhibited adequate antibacterial activity, and all showed no microorganism growth except for gentamicin in adsorbent-free BACT/ALERT SA media. The overall recovery rate of FA Plus media was 87.5% (35/40), which was higher than that of DL aerobic medium (42.9%, 15/35; *p* < 0.0001) and Thermo Aerobic media (12.5%, 5/40; *p* < 0.0001). In anaerobic culture, the overall recovery rate of FN Plus media was 97.4% (32/35), which was higher than that of 2.9% (1/35) of DL media (*p* < 0.0001) and 14.3% (5/35) of Thermo media (*p* < 0.0001; Table 2, Fig. 1a). The neutralizing effects of the antimicrobial agents in aerobic and anaerobic media from the same manufacturer were consistent, except that the recovery rate of the DL aerobic media was higher than that of the anaerobic media (42.9% vs. 2.9%, *p* < 0.0001; Table 2, Fig. 1b).

**Table 2.** Summary of recovery rate (%) in three different culture media (BacT/Alert FA/FN Plus media, DL media,Thermo media) containing antimicrobials by ATCC strains.

| ATCC strains | Recovery rate(%) |  |  |  |  |  |  |  |  |
| --- | --- | --- | --- | --- | --- | --- | --- | --- | --- |
|  | BACT/ALE RT FA Plus | BACT/ALE RT FN Plus | As a pair | DL Aerobic | DL Anaerobic | As a pair | Thermo Aerobic | Thermo Anaerobic | As a pair |
| *S.pneumonia* | 100.0(5/5) | 100.0(5/5) | 100.0(5/5) | 0.0(0/5) | 0.0(0/5) | 0.0(0/5) | 0.0(0/5) | 0.0(0/5) | 0.0(0/5) |
| *P.aeruginosa* | 100.0(5/5) | -<sup>a</sup> | - | 100.0(5/5) | - | - | 0.0(0/5) | - | - |
| *B. fragilis* | - | 40.0(2/5) | - | - | 0.0(0/5) | - | - | 0.0(0/5) | - |
| *E. coli* | 50.0(5/10) | 100.0(10/10) | 50.0(5/10) | 50.0(5/10) | 0.0(0/10) | 0.0(0/10) | 50.0(5/10) | 50.0(5/10) | 50.0(5/10) |
| *S.aureus* | 100.0(15/15) | 100.0(15/15) | 100.0(15/15) | 33.3(5/15) | 6.7(1/15) | 6.7(1/15) | 0.0(0/15) | 0.0(0/15) | 0.0(0/15) |
| *C.albicans* | 100.0(5/5) | - | - | - | - | - | 0.0(0/5) | - | - |
| Total | 87.5(35/40) | 91.4(32/35) | 83.3(25/30) | 42.9(15/35) | 2.9 ( 1/35 ) | 3.3(1/30) | 12.5(5/40) | 14.3(5/35) | 16.7(5/30) |

**Notes:** <sup>a</sup> Lack of data because the microorganism was cultured only in aerobic or anaerobic media.

**FIG 1:**
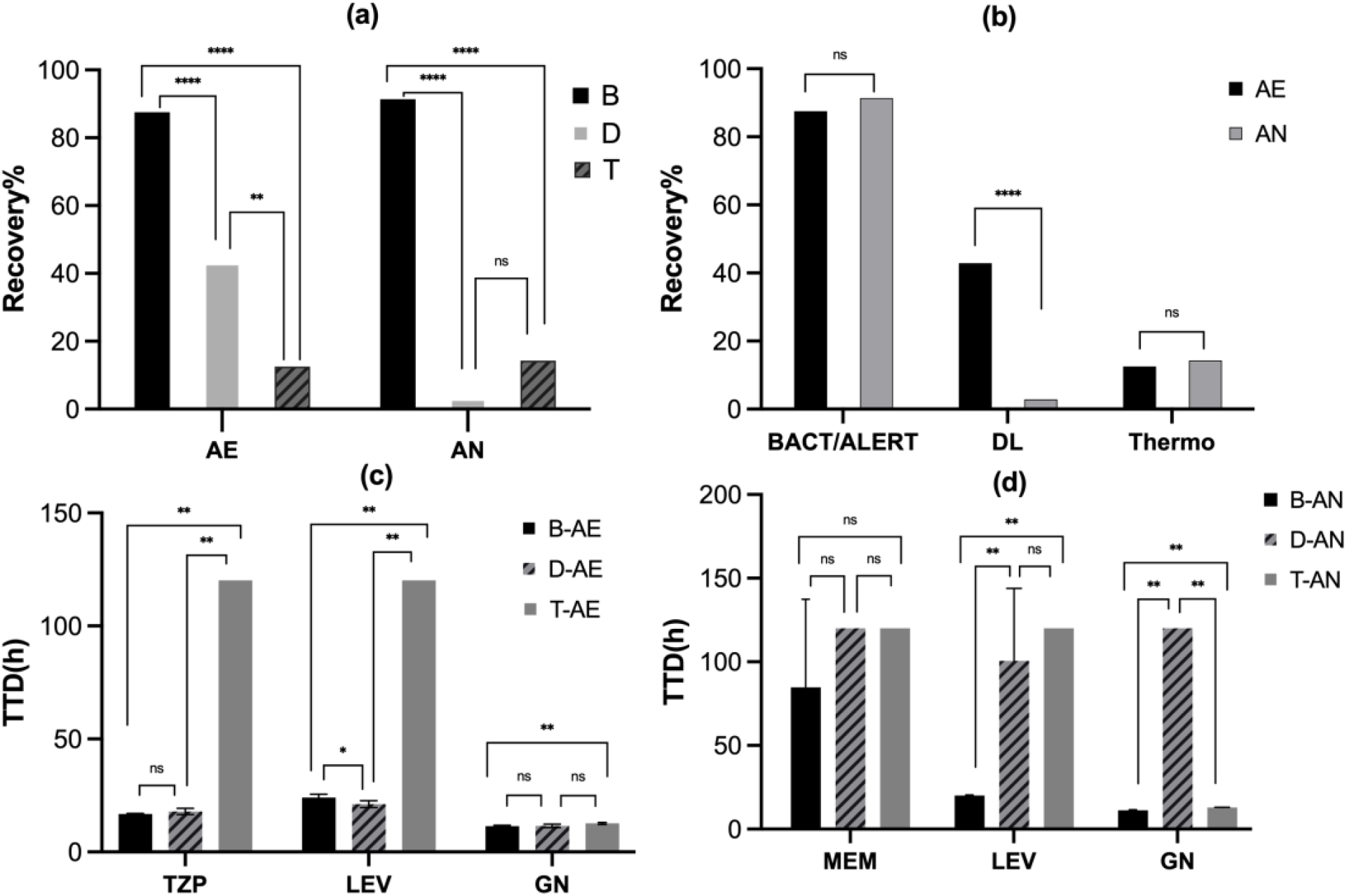
**(a)** Recovery rate of microorganisms in Aerobic (BacT/ALERT Plus, DL and Thermo) or anaerobic (BacT/ALERT Plus, DL and Thermo) culture media containing antimicrobials. **(b)** Recovery rate of microorganisms in different (aerobic and anaerobic) culture media containing antimicrobials. **(c)** Time to detection (TTD) of microorganisms recovered from BACT/ALERT FA Plus, DL aerobic and Thermo aerobic culture media containing different antimicrobials (piperacillin/Tazobactam (TZP), levofloxacin (LEV), and gentamicin (GN). **(d)** Time to detection (TTD) of microorganisms recovered from BACT/ALERT FN Plus, DL anaerobic, and Thermo anaerobic culture media containing different antimicrobials (meropenem (MEM), levofloxacin (LEV), and gentamicin (GN)). **Abbreviations: B**, BACT/ALERT culture (FA Plus and FN Plus) media; **D**, DL culture media; **T**, Thermo culture media; **AE**, aerobic media; **AN**, anaerobic media; **ns**, not significant; **\***, *P* < 0.05; **\*\***, *P* < 0.01; **\*\*\***, *P* < 0.001; **\*\*\*\***, *P* < 0.0001. Error bars indicate standard deviations (SD).

Gentamicin demonstrated the highest recovery rate as nearly all the tested strains were recovered in both culture media except for the DL anaerobic bottle, and the BACT/ALERT (FA plus and FN Plus) culture media showed a slightly shorter TTD of gentamicin than Thermo culture media (*p* < 0.01; Table 3, Fig. 1c, Fig. 1d).

**Table 3.**
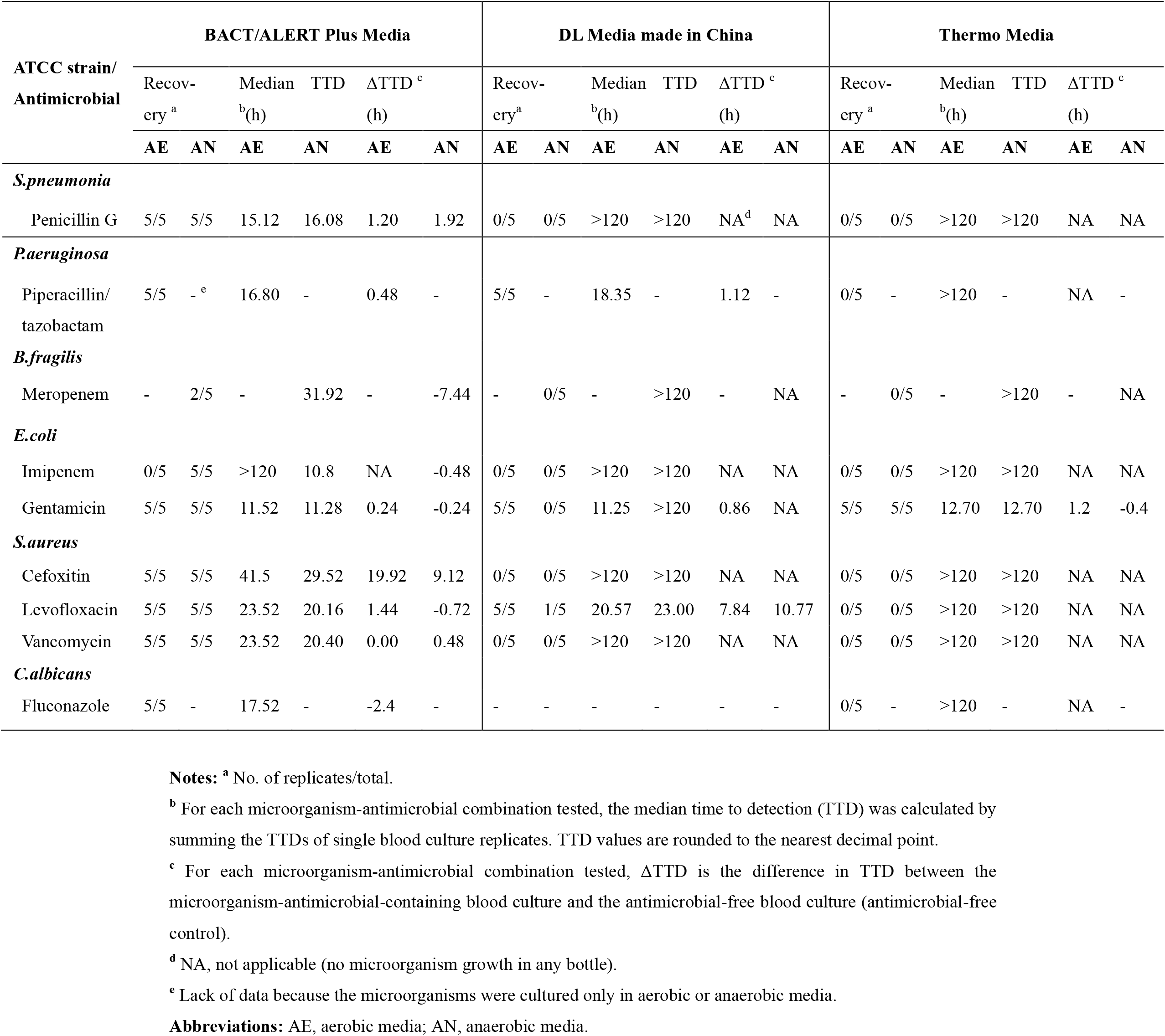
Time to detection of microorganisms recovered from three different aerobic and anaerobic media at %PSL concentration tested in blood culture quintuplicates.

Piperacillin-tazobactam and levofloxacin achieved the highest recovery rate, with all the test strains being recovered in BACT/ALERT Plus media and DL media. TTD of piperacillin-tazobactam in aerobic atmosphere showed no significant difference between the two culture media; BACT/ALERT FN Plus culture showed slightly shorter TTD (−2.8 h) of levofloxacin than DL anaerobic culture media (*p* < 0.008), whereas BACT/ALERT FA Plus culture showed slightly longer TTD (3.0 h) of levofloxacin than DL aerobic culture media (*p* < 0.008; Table 2, Figs. 1c-d). Furthermore, only the BACT/ALERT system revealed 100% recovery with approximately all the tested antimicrobial agents, except carbapenems, which were significantly higher than those of the DL system (penicillin G, *p* < 0.0001; cefoxitin, *p* < 0.0001; vancomycin, *p* < 0.0001; Table 3). For carbapenems, meropenem exhibited the lowest recovery rate in all blood cultures (13.3%, 2/15), and most of the microorganisms were only recovered in BACT/ALERT FN Plus media (40.0% vs. 0.0% vs 0.0%, *p* = 0.44). Similar to imipenem, with a low recovery rate in all blood cultures (16.7%, 5/30), microorganisms were only recovered in BACT/ALERT FN Plus media (50.0% vs. 0.0% vs. 0.0%, *p* = 0.03); For fluconazole, microorganisms were only recovered from BACT/ALERT FN Plus media.

Our research also focused on comparing TTD between BACT/ALERT Plus media and DL media, as Thermo blood culture media only exhibited the recovery of organisms with gentamicin. Seven microorganism-antimicrobial combinations were identified as effective in recovering organisms in both aerobic culture systems. Moreover, shorter TTD in FA Plus culture media than in DL aerobic culture media was observed in four (57.1%) combinations. In two anaerobic culture systems, seven microorganism-antimicrobial combinations were detected to recover organisms, with all seven (100.0%) indicating shorter TTD in FN Plus media than in the corresponding DL anaerobic media (Table 3). Furthermore, no statistically significant difference in TTD between FA Plus and DL aerobic media (*p* = 0.350) was observed.

Table 3 summarizes the differences in TTDs between media spiked with microorganism-antimicrobial combinations and antimicrobial-free media (ΔTTD) (7). For BACT/ALERT (FA Plus and FN Plus) media, the highest ΔTTD (19.9 h aerobic, 9.1 h anaerobic) was detected for cefoxitin-*S. aureu*s, whereas for DL media, the ΔTTD of levofloxacin-*S. aureus* was also high (7.8 h aerobic, 10.8 h anaerobic). Negative ΔTTDs were observed in some microorganism-antimicrobial combinations, particularly in fluconazole-*C. albicans* (−2.4 h) in BACT/ALERT FA Plus media and meropenem-*B. fragilis* (−7.4 h) in BACT/ALERT FN Plus media.

## Discussion

Numerous techniques have been developed to neutralize antimicrobials in blood culture media; each has advantages and disadvantages (3–4). BACT/ALERT (FA Plus and FN Plus) media containing improved ZLAPB resin adsorption has three distinct principles: synergistic ion exchange, van der Waals forces, and covalent bonds, which optimize the adsorption performance of antimicrobials, including cephalosporins, imipenem, meropenem, daptomycin, cephalosporin, carbendazim, and other high-order antimicrobial agents. Compared with the single adsorption principle of activated charcoal and common resins, culture media containing APB resin and ZL covalent bond additives can adequately achieve the neutralization requirements of antimicrobials without the interference of trace elements and nutrients necessary for bacterial growth (8–13). We observed significantly higher microorganism recovery rates with the BACT/ALERT Plus media than the other two culture media (DL and Thermo) containing antimicrobials both in aerobic and anaerobic cultures. Similarly, there was a higher detection rate for microorganisms in at least one bottle of each pair. However, the BACT/ALERT system exhibited a more satisfactory detection rate and higher recovery rates than those reported elsewhere (5–7), which was probably due to the larger inoculum (30-10^2^ CFU/bottle), appropriate inoculation volume (approximately 10 mL), and susceptible ATCC strains used in our research. Almost all antibacterial categories were successfully neutralized in both BACT/ALERT (FA Plus and FN Plus) culture media, except for carbapenems. For carbapenems, microorganisms were only recovered in BACT/ALERT FN Plus media. This outcome was expected to be affected by carbapenem neutralization of BACT/ALERT FN Plus media relying on the adsorption of ZL cysteine covalent bonds (7–10), and cysteine is essential for anaerobic environments. Similar to previous studies concerning BACT/ALERT (FA Plus and FN Plus) culture media, microbial-imipenem and microbial-meropenem detection rates were lower, 5/10 (50.0%) and 2/5 (40%), respectively (5–7). Our result indicated the lowest recovery rate of *B. fragilis* with meropenem in BACT/ALERT blood culture, possibly due to the low recovery rate of *B. fragilis* with meropenem. A potential explanation for this is the low MIC and high Cmax/MIC quotient for the strain-antimicrobial combination. The new *B. fragilis*-meropenem recovery combination can be improved by diluting the concentration of meropenem or increasing the quantity of the novel ZLAPB resin. We also observed that the concentration of free levofloxacin quickly fell below the strain’s MIC range in BACT/ALERT FA Plus media within 10 min of incubation [14]. Hence, to prevent the majority of antimicrobials from being absorbed, a bacterial suspension was added before the antimicrobial solution. We achieved a more consistent recovery rate of the *S. aureus-levofloxacin* combination in BACT/ALERT (FA Plus and FN Plus) culture media compared to previous investigations (6,14).

The anaerobic media comprising distinct media and *Candida spp*. being facultative anaerobes with extended generation times under anaerobic conditions could influence the recovery of numerous pathogens, particularly when exposed to antimicrobial agents. Therefore, we excluded *C. albicans* fluconazole recovery in anaerobic media (15). For the domestically manufactured DL culture media with common resin, we further excluded *C. albicans*-fluconazole recovery in aerobic media as the positive controls were false, and *C. albicans* cannot recover from DL culture media of free fluconazole. DL aerobic culture media with common resin demonstrated better and higher recovery rates of piperacillin-tazobactam and levofloxacin than DL anaerobic and Thermo culture media. A significant difference (*P* < 0.0001) in antimicrobial neutralization was observed between the DL aerobic and anaerobic media due to the variation in the oxygen consumption niche of spiked microorganisms.

The overall performance of the Thermo culture media without any adsorbent was inferior to that of the other two media owing to lower neutralizing activity, relying on an optimal 1:9 blood-broth dilution to neutralize antimicrobial agents. However, the observations indicated a high recovery rate of the microorganism-gentamicin combination in all three media as gentamicin is probably easily affected by oxygen consumption, SPS anticoagulant, antimicrobial adsorbent, and dilution.

TDD is a significant parameter to evaluate the performance of commercial automatic blood culture systems [5–6,16]. Generally, BACT/ALERT FN Plus media demonstrated shorter TDDs than DL anaerobic media because of low microbial recovery rates. Moreover, there was no statistically significant difference in TTDs between FA Plus media and DL aerobic media. The BACT/ALERT FA Plus media did not outperform DL aerobic media in terms of TDDs, given that their microbial recovery rates were nearly equivalent. For gentamicin, BACT/ALERT (FA plus and FN Plus) culture media showed a slightly shorter TTD than the Thermo culture media. However, the potential impact of the typical several-hour differences on decision-making in clinical settings remains unclear. Additional prospective clinical research is required to evaluate the influence of TTD on the initiation or change in antibiotic treatment strategies.

Regarding ΔTTD, another crucial parameter to assess the antimicrobial inactivation ability of the examined automatic blood culture media [7], delays of at least 3 h compared with positive results in control (antimicrobial-free) blood culture media were only observed for cefoxitin-*S. aureu*s in the BACT/ALERT (FA Plus and FN Plus) culture media pair and levofloxacin-*S. aureus in the* DL culture media pair. Negative Δ TTDs were also observed in certain microorganism-antimicrobial combinations, specifically in fluconazole-*C. albicans* (−2.4 h) in BACT/ALERT FA Plus media, consistent with previous reports [6]. Moreover, negative ΔTTDs for meropenem-*B. fragilis* (−7.4 h) in BACT/ALERT and FN Plus media were expected to be affected by the novel resin ZLAPB. Furthermore, delayed ΔTTDs were more frequently observed in aerobic media. Consequently, the outcome of antimicrobial inactivation may be influenced to some extent by the type of antimicrobials and/or microbe species, as well as the interaction between the microbial and a specific antimicrobial in complicated environments, including blood culture media.

This study has some limitations. First, horse blood was used for the simulation instead of human blood. Second, the evaluation was limited to susceptible ATCC strains, excluding resistant clinical isolates. Finally, only a restricted number of bacterial and antimicrobial species were assessed, and those frequently isolated in a clinical setting, including MRSA, *enterococci*, and filamentous fungi, were not examined. Additional studies, including a wider variety of microbial and antimicrobial species, will yield further insights into the comparative assessment of blood culture media for antimicrobial neutralization.

## Conclusion

Absolute advantages were revealed in our simulated study using BACT/ALERT (FA Plus and FN Plus) media containing the novel resin. Except for carbapenems, most of the tested broad-spectrum antimicrobial agents under clinically significant concentrations can be effectively neutralized. To optimize recovery chances for patients who have previously undergone antimicrobial therapy, relatively more expensive media (FA Plus and FN Plus) can be selectively employed. The results indicated that the DL aerobic culture medium with common resin was significantly more efficient in recovering challenge organisms with piperacillin/tazobactam and levofloxacin than adsorbent-free Thermo culture media.

Emphasis should be placed on enhancing the neutralization of carbapenems with low efficiency in all three media. Further comparative studies using blood samples from patients undergoing antimicrobial therapy will provide additional information on the performance of culture systems for antimicrobial neutralization.

## Acknowledgments

This work was supported by the Natural Science Foundation of Jiangxi Province (No. 20202BAB216021), the Health Commission of Jiangxi Province (No. 20201034). Youth Science Fund of the Science and Technology Program of 2^nd^ affiliated Hospital, Jiangxi Medical College, Nanchang University(2019YNQN12005)

